# Metabolic profiling suggests two sources of organic matter shape microbial activity, but not community composition, in New Zealand fjords

**DOI:** 10.1101/745273

**Authors:** Sven P. Tobias-Hünefeldt, Stephen R. Wing, Federico Baltar, Sergio E. Morales

## Abstract

Fjords are semi-enclosed marine systems with unique physical conditions that influence microbial communities structure. Pronounced organic matter and physical condition gradients within fjords provide a natural laboratory for the study of changes in microbial phylogeny and metabolic potential in response to environmental conditions (e.g. depth). In the open ocean new production from photosynthesis supplies organic matter to deeper aphotic layers, sustaining microbial activity. We measured the metabolic diversity and activity of microbial communities in fjords to determine patterns in metabolic potential across and within fjords, and whether these patterns could be explained by community composition modifications. We demonstrated that metabolic potential and activity are shaped by similar parameters as total (prokaryotic and eukaryotic) microbial communities. However, we identified increases in metabolic diversity and potential (but not in community composition) at near bottom (aphotic) sites consistent with the influence of sediments in deeper waters. Thus, while composition and function of the microbial community in the upper water column was likely shaped by marine snow and sinking POM generated by new production, deeper sites were strongly influenced by sediment resuspension of benthic organic matter generated from this or other sources (terrestrial, chemoautotrophic, microbial carbon loop), uncoupling the community composition and function dynamics.

## Introduction

Fjords are unique environments, representing modified marine ecosystems mixing freshwater, terrestrial and marine inputs. As such, influences on structure and function of microbial communities are linked to changes in environmental conditions associated with each input, including alternate organic carbon sources (e.g. tannins, terrestrial, marine and freshwater sources), as well as modified salinity, nutrient, and light environments (Mckee et al., 2002; Pulchan et al., 2003; Cui et al., 2016). Moreover, due to these strong environmental gradients, fjords are ideal natural laboratories to study marine microbial communities and the controls of their phylogenetic and functional diversity. However, the energy sources supporting primary production and heterotrophic activity in fjords, and how they change in correlation to observed community changes, remain poorly defined. In open ocean systems primary productivity by surface phytoplankton mediates the downward flux of particulate carbon, transferring energy to aphotic zones. This unidirectional transfer of carbon through microbial/biological biomass from surface waters to deeper layers is termed the biological carbon pump (Jiao et al., 2010; Jiao and Zheng, 2011; Legendre et al., 2015). This process is also expected to dominate in fjords where carbon is predominantly linked to phytoplankton production (Albright et al., 1986; Amy et al., 1987; Alldredge et al., 2002), sustaining a significant portion of heterotrophic respiration (Iturriaga and Hoppe, 1977). Nevertheless, fjord benthic community studies have demonstrated that microbial reworking of refractory organic matter from terrestrial sources is an additional important source of carbon to deep communities (McLeod and Wing 2007, McLeod and Wing 2009, McLeod Wing and Skilton 2010). Despite this, we lack an integrated view of microbial metabolic potential across fjords and specific information about microbial populations possibly linked to them, providing a mechanistic understanding of their selection. This limits our understanding of how this ecosystem is sustained and shaped.

We previously examined the patterns in microbial community composition relative to variability in environmental factors among fjords in the New Zealand Fiordland system (Tobias-Hünefeldt et al., 2019). But links between patterns in phylogenetic and functional diversity in these fjords remained unaddressed. In the present study we utilised functional potential profiling (via Biolog Ecoplates), bacterial abundance, heterotrophic production (via ^3^H-leucine incorporation) and prokaryotic/eukaryotic community composition (via 16S and 18S rRNA amplicon sequencing) to compare community metabolic diversity and potential, and how it related to known drivers of microbial community changes across six different fjords in New Zealand. We found that community metabolic potential and diversity at surface sites follow similar patterns to those observed when examining whole community composition. However, a high resolution analysis along a depth profile of a fjord indicates two potential drivers of metabolic diversity and potential (i.e. vertical transfer of carbon via suspended particulate organic matter through the biological carbon pump, and resuspension of organic matter from sediments linked to the benthic microbial loop and terrestrial carbon sources).

## Results and Discussion

The present study was carried out in six fjords within New Zealand’s Fiordland system, specifically Breaksea Sound, Chalky Inlet, Doubtful Sound, Dusky Sound, Long Sound, and Wet Jacket Arm, utilising an identical sampling scheme as described in (Tobias-Hünefeldt et al., 2019). Analyses were divided into three categories, a multi-fjord analysis comprising five of the tested fjords (excluding Long Sound), a high resolution study along Long Sound’s horizontal axis, and a depth profile of Long Sound’s deepest location. Total community composition (via 16S and 18S sequencing) and metabolic potential did not significantly covary across the five studied fjords (Mantel, r = <0.01, p = 0.47), Long Sound’s horizontal transect (Mantel, r < 0.01, p = >0.05), or Long Sound’s depth profile (Mantel, r = <0.22, p = >0.05). However, depth covaried with both metabolic potential and community structure among all fjords, across the horizontal transect at Long Sound, and along Long Sound’s depth profile (Figure 1, Figure S1-S3, Table S1). Significant differences in metabolic potential with depth were observed both across multiple fjords (Anosim: R= 0.10, P value= 0.03) and along a transect from the entrance of the ocean to the head of Long Sound (Anosim: R= 0.27, P value= <0.01) (Figure 1). Microbial community changes along the horizontal axis were stronger between surface and 10 m communities (Mantel, Multifjord – r = 0.21, p = <0.01, Transect – prokaryotes r = 0.47, p = <0.01, eukaryotes r = 0.56, p = <0.01), as opposed to horizontal location (Mantel, Multifjord – r = 0.08, p = 0.04, Transect – prokaryotes r = 0.21, p = 0.01, eukaryotes r = 0.13, p = 0.07) (Figure S2-3).

**Figure 1.**
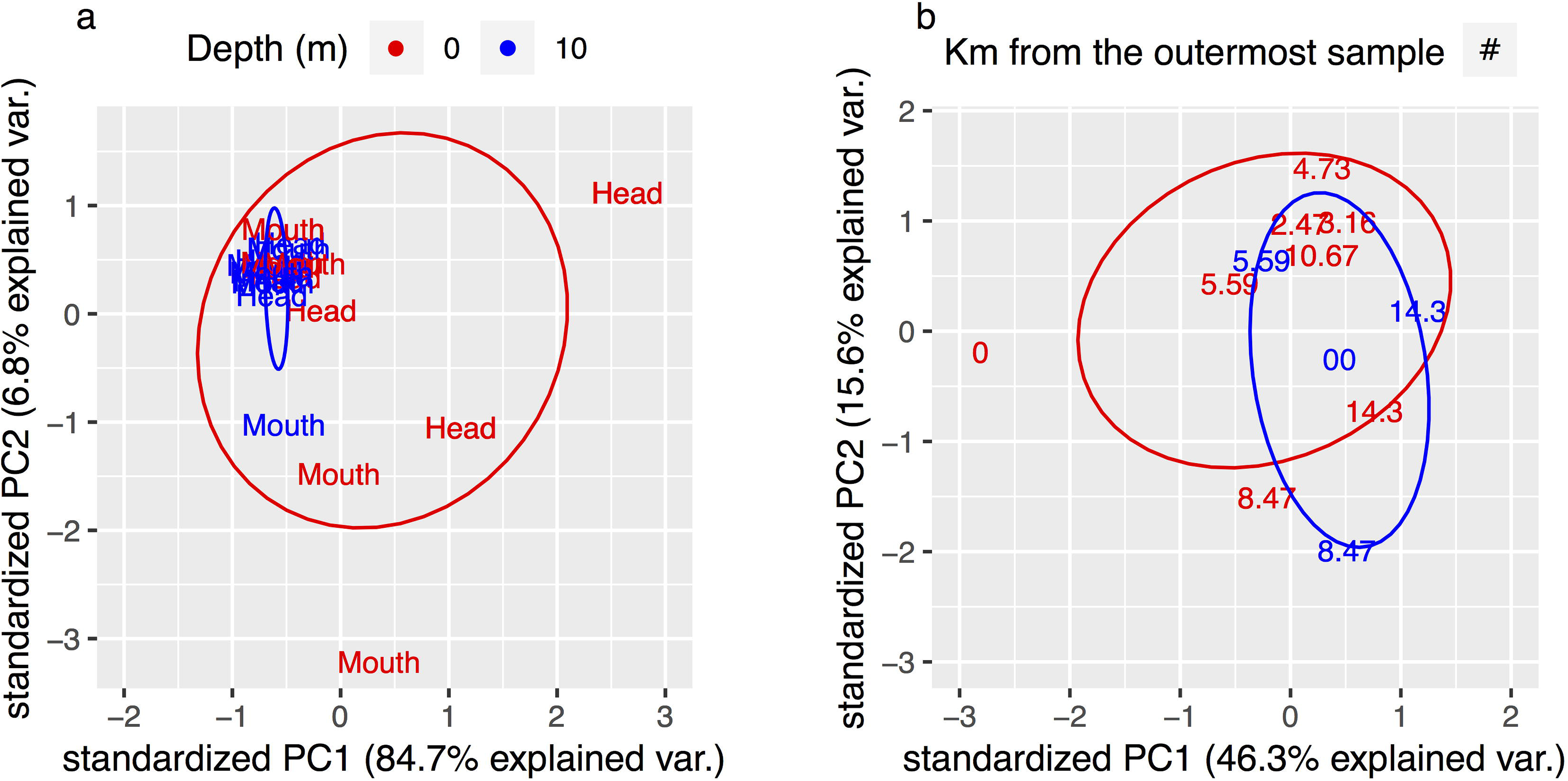
Biolog Ecoplate derived surface vs. 10 m PCA. Depth separated samples for Multifjord data (a) Long Sound’s horizontal transect (b) calculated into a PCA plot. Text labels representing the horizontal location; either near the head/mouth of the fjord (a), or being defined by the Km from the outermost sample (b). Ellipses represent the 95% confidence interval. 150 μL of sample were utilised per Biolog Ecoplate well, and then incubated for 7 days at 4°C. Colour patterns assessed at OD A590 nm.

Across the five fjords (excluding Long Sound), surface samples displayed higher metabolic potential (i.e., average metabolic rate [AMR]) compared to 10 m samples (Wilcox test, W = 425, p = <0.01) with samples from the fjord’s head having a higher rate in general (Wilcox test, W = 0, p = <0.01). Horizontal sampling location affected the observed variance, with higher variability in metabolic potential observed in samples collected near the fjord entrance (Figure 1 b, Table S1). While activity was not consistent along the length of the longest fjord, a sustained elevated activity was seen at surface compared to 10 m depths (Figure S4). Heterotrophic production (via leucine incorporation) was not significantly correlated with microbial abundance within the five studied fjords and Long Sounds horizontal axis (Mantel – Multifjord r = 0.04, p = 0.22, Horizontal r = 0.04, p = 0.32), consistent with either differences in grazing pressure between locations or a small proportion of cells driving a large portion of the productivity. Along the depth profile, prokaryotic abundance and production were significantly correlated (Mantel, r = 0.60, p = 0.01); however clear differences were present, such as a more gradual difference in abundance compared to the large drop in productivity from the surface to 10 m.

To further explore these depth linked changes we focused on a high resolution depth profile of the deepest fjord. We hypothesized that metabolic rate and diversity would be driven by marine snow linked to photosynthetic primary producers at the surface (e.g. phytoplankton and macroalgae) (Figure 2a) leading to a steady decrease in metabolic potential as resources were depleted with increases in depth. Any deviation altering the slow loss of metabolic potential would be linked to extraneous sources of nutrients uncoupled from surface activity. We observed a steady loss of metabolic diversity, and rate, from surface to 100 meters (Figure 2 b, c), with a sustained increase past this point. The observed pattern was consistent with measurements for bacterial production, but not abundance, that decreased continuously until reaching equilibrium from 200 m onwards (Figure 2 d). These changes were associated with shifts in specific carbon utilization potential, where carbohydrate metabolism decreased from 12.7% to 6.8%, as carboxylic acid utilization increased from 12.0% to 29.5% (Figure 2 e). This likely reflected the diminishing abundance of readily mineralizable substrates with depth, and the increase in recalcitrant sources of carbon and energy. Consistently, we also observed increases in phosphorylated chemical metabolism peaking at 40 and 360 m (Figure 2e) as expected from utilization of phosphorous at the surface during blooms (Tiselius and Kuylenstierna, 1996). However, observed changes in metabolic potential did not reflect changes in prokaryotic or eukaryotic community composition, suggesting that while the community members were relatively consistent past a certain depth (10 m for eukaryotes and 40 m for prokaryotes) functional potential changed dynamically past 100 m, regaining metabolic potential with proximity to the bottom (Figure 2 f) to a point closely resembling the metabolically active surface.

**Figure 2.**
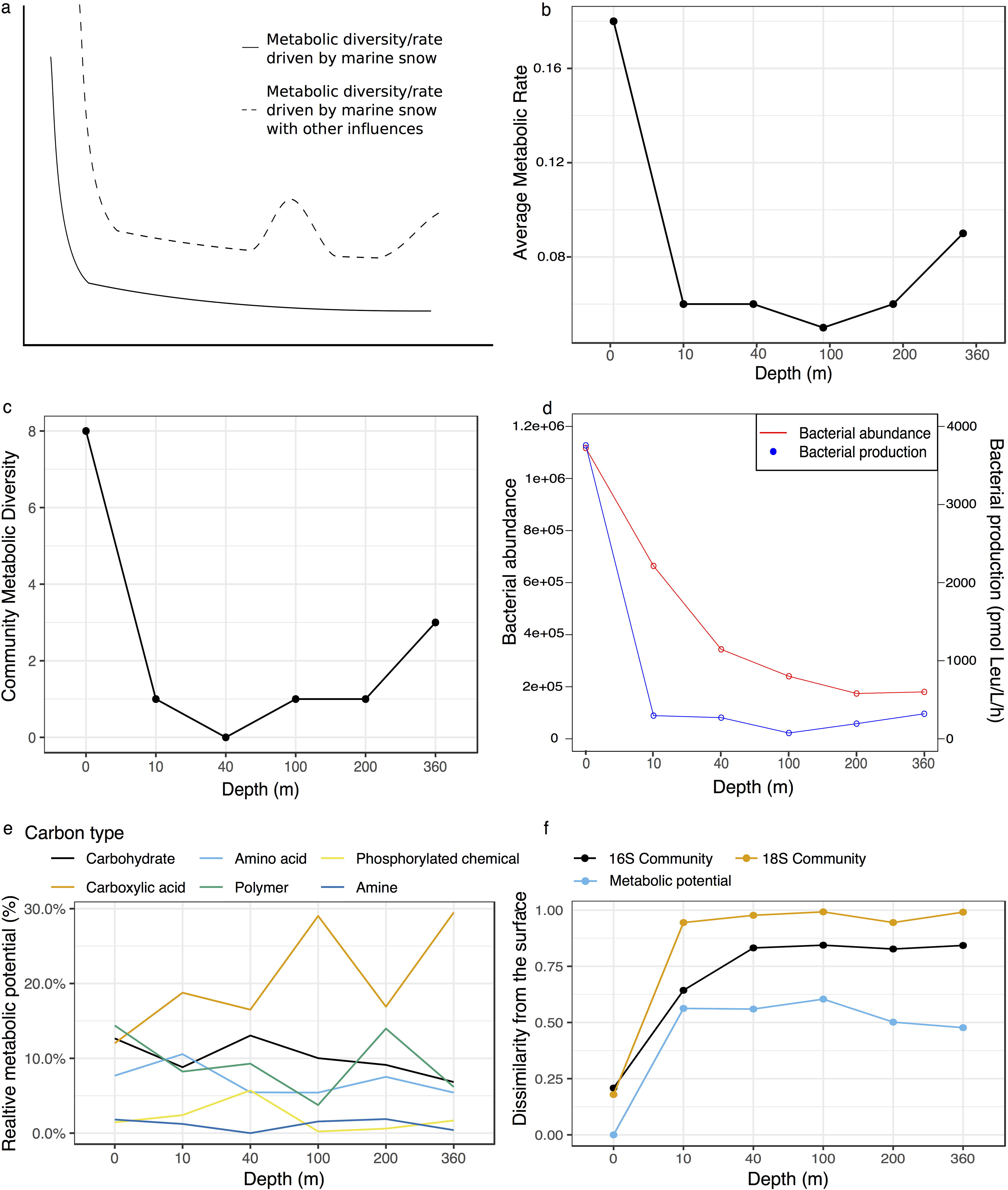
Benthic and surface influence on metabolic potential. A model of purely surface vs. surface and benthic influences is shown (a), the Biolog Ecoplate plate derived Average Metabolic Rate (AMR, b), Community Metabolic Diversity (c), and relative metabolic potential (e). Bacterial abundance and productivity (c), and taxonomic and Biolog plate derived dissimilarity from the surface (f). Different carbon source groups were displayed in various colours (carbohydrates are blue, carboxylic acids are orange, amino acids are light blue, polymers are green, phosphorylated chemicals are yellow, amines are dark blue), as well as the Bray-Curtis dissimilarity measures (16S community being black, 18S community being orange, and Biolog derived metabolic potential being light blue). Communities were sequenced using the Earth Microbiome protocol (Thompson et al., 2017), OTUs were generated using QIIME (Caporaso et al., 2012), UCLUST (Edgar, 2010) and SILVA (Quast et al 2013).

Our results demonstrate that while metabolic potential and activity in fjords is linked to similar parameters as microbial community composition across surface or near surface sites, distinct selective pressures exists at aphotic sites, ultimately affecting the link between phylogenetic and metabolic diversity. The observed pattern is contrary to our initial hypothesis and demonstrates that additional refractory sources of organic matter including resuspension of terrestrial organic matter associated with benthic communities are important contributors to microbial activity in fjords. We propose that this reflects the influence of the benthic microbial loop and incorporation and breakdown of terrestrial organic matter in fjordic sediments. Sediment resuspension through either wave or wind action (Pickrill, 1987; Christiansen et al., 1992), as well as advection and biological activity provides mechanisms for increased availability to the microbial community in the deep water column. Sediment resuspension is known to increase metabolic activity of microbes (Flindt and Kamp-Nielsen, 2017) which is likely an important process in this deep water site where loose sediments are organically rich reflecting suspended particulate matter [large amounts of fibrous woody material, finer indeterminate organic plankton, faecal pellets with a small terrestrial influence] (Pickrill, 1987; Pickrill, 1993). The observed pattern suggests that resuspension could also be driven by bottom feeding organisms that can resuspend sediments, increasing suspended carbon and its utilization in near bottom sites (Yahel et al., 2008), influencing the relation between marine diversity and the metabolic potential of marine microbes.

## Acknowledgements

We thank the officers and crew of *RV Polaris II* and science staff involved. We also thank M. Meyers, O. Shatova, B. Dagg and J. Wenley for their contribution to data collection. FB was supported by a Rutherford Discovery Fellowship (Royal Society of New Zealand). Field sampling was supported by a Royal Society of New Zealand Marsden Grant (UOO1008) to SW.

## Data availability

The sequence data from this study have been deposited in NCBI under BioProject PRJNA540153. All data generated and/or analysed during the study is available within the GitHub repository, https://github.com/SvenTobias-Hunefeldt/Fiordland_2019b/.

Supplementary information is available at Frontiers in Marine Science Journal’s website’.

---

**Figure S1. Microbial beta-diversity of five fjords.** The fjord of origin, sample region, and depth were used to identify sample origin. Dissimilarity was assessed using Bray-Curtis distance matrices based on OTUs at 97% similarity.

**Figure S2. Prokaryotic beta-diversity across Long Sound.** Beta-diversity based on 16S data for Long Sound’s horizontal axis. Dissimilarity was assessed using Bray-Curtis distance matrices based on OTUs at 97% similarity.

**Figure S3. Eukaryotic beta-diversity across Long Sound.** Beta-diversity based 18S data for Long Sound’s horizontal axis. Dissimilarity was assessed using Bray-Curtis distance matrices based on OTUs at 97% similarity.

**Figure S4. Average Metabolic rate across Long Sound.** The average metabolic rate (AMR) across Long Sound’s horizontal axis, colour separating depth (red being surface and blue 10 m).

