## Supplementary figures and images for "Metabolic profiling suggests two sources of organic matter shape microbial activity, but not community composition, in New Zealand fjords"

### Supplemental Figure S1

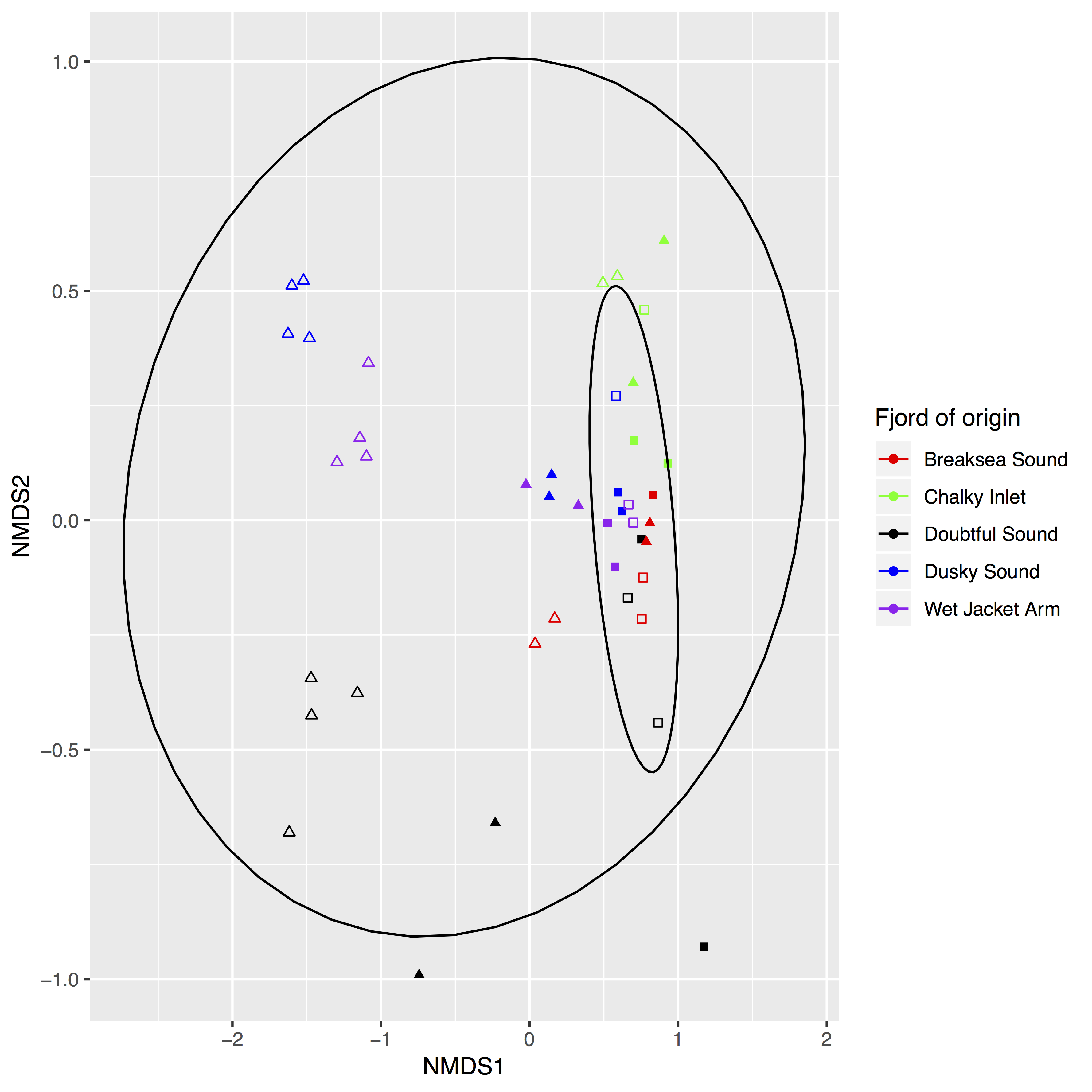

### Supplemental Figure S2

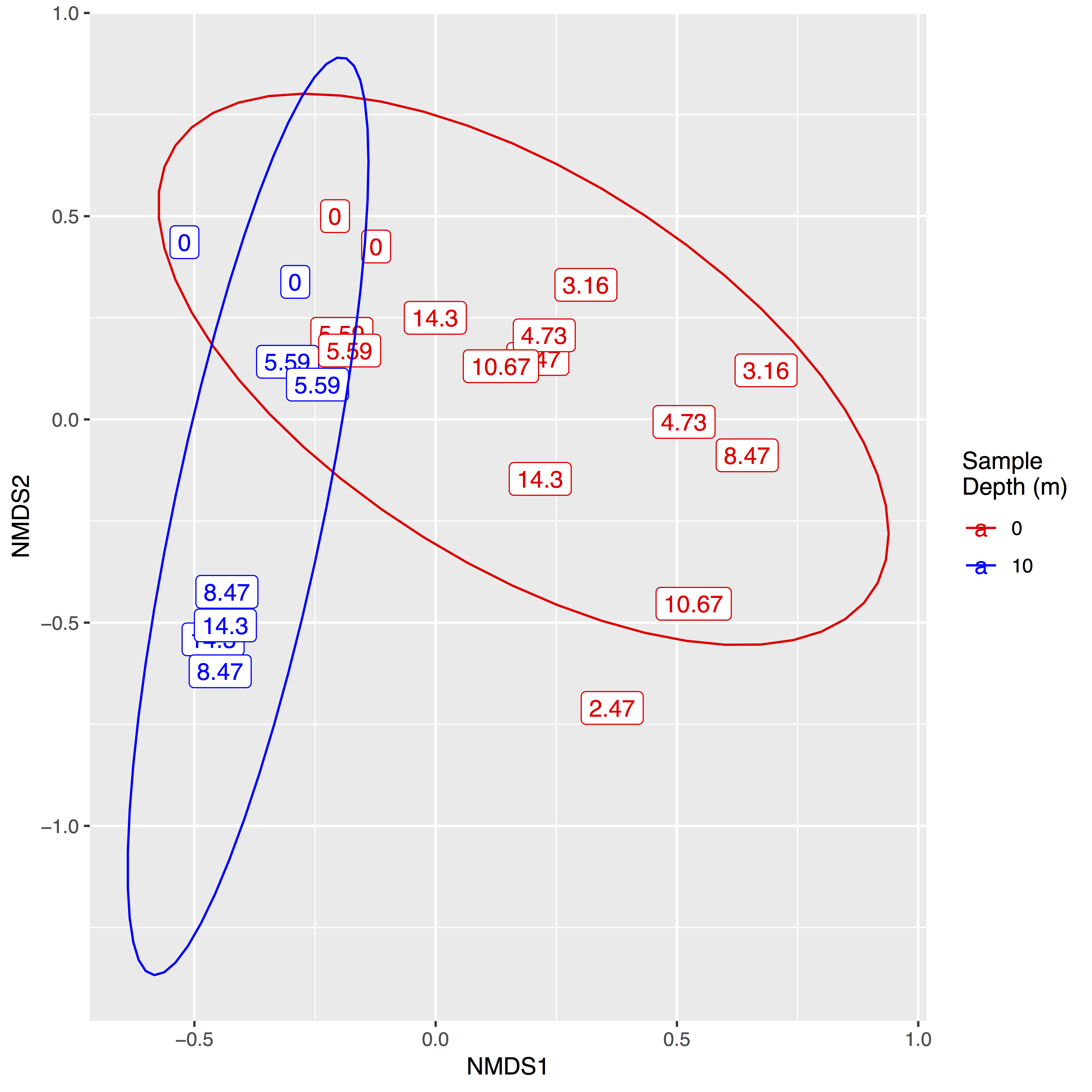

### Supplemental Figure S3

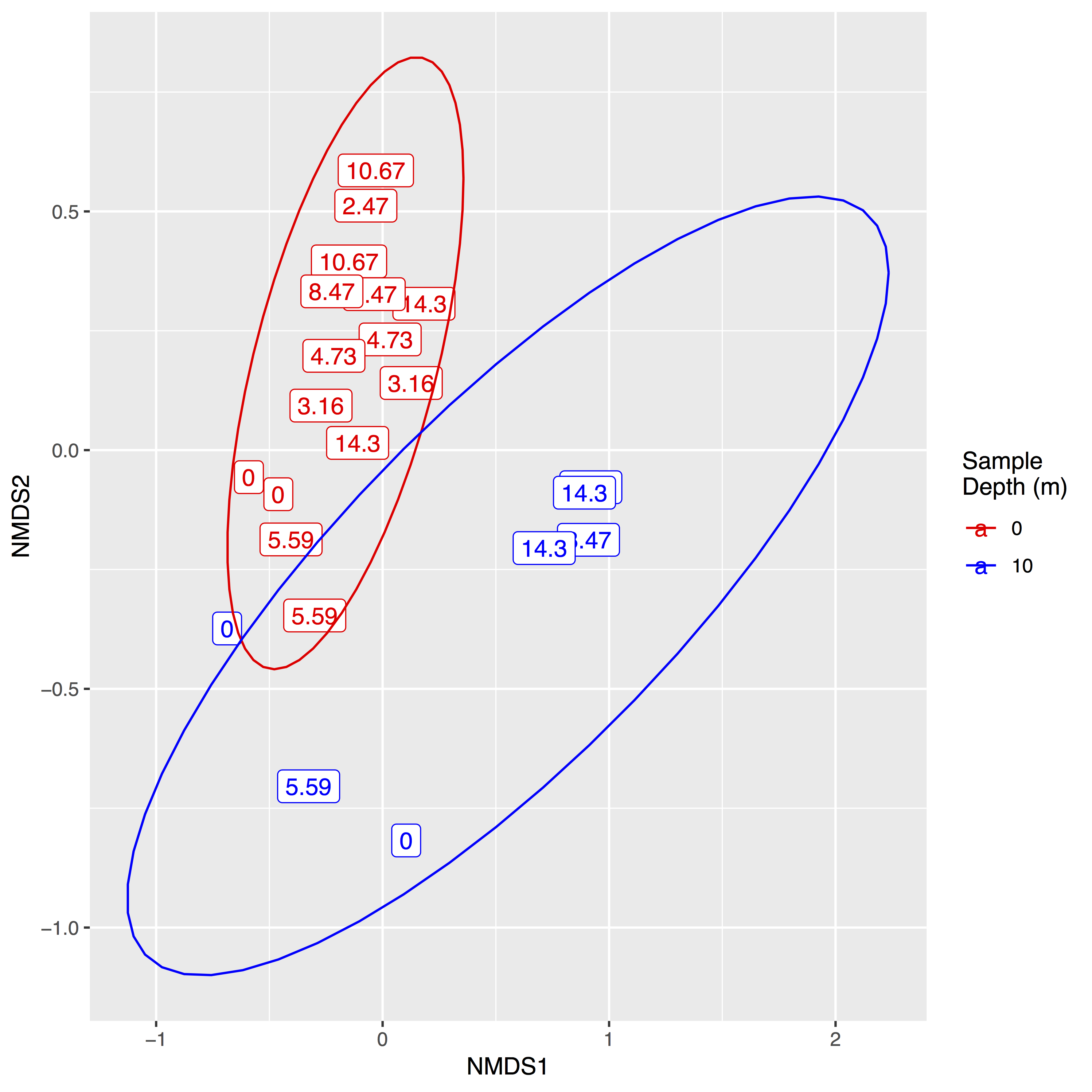

### Supplemental Figure S4

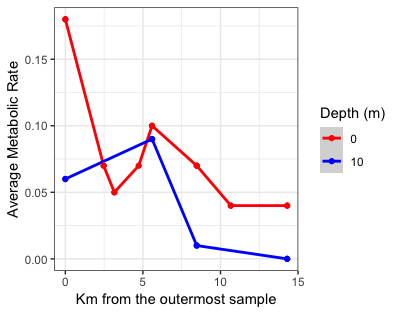
